# Intra-Abdominal Bowel Dilation in Experimental Gastroschisis is Associated with a Modifiable Transcriptomic Program of Intestinal Dysfunction

**DOI:** 10.64898/2026.05.29.728798

**Authors:** Mary Elizabeth Guerra, Tomohiro Arai, Luc Joyeux, Casey C. Baxter, Sourav Bose, Shiyanth Thevasagayampillai, Hui Li, Ling Yu, Varun Akondy, Marianna Scuglia, David Basurto, Emma Van den Eede, Simen Vergote, Kanokwaroon Watananirum, Wasinee Tianthong, Francesca Russo, Paolo De Coppi, Preethi H. Gunaratne, Lily S. Cheng, Michael A. Belfort, Swathi Balaji, Jan Deprest, Sundeep G. Keswani

**Author notes:** Co-first author. Co-last author (Jan Deprest, Sundeep G. Keswani). Corresponding Author: Sundeep G. Keswani, MD, MBA, FACS, 6701 Fannin St, Houston, TX 77030, Jan Deprest MD, PhD.

## Abstract

**Objective:** To characterize intestinal transcriptional profiles in gastroschisis, their temporal evolution, and response to fetal intervention.

**Summary Background Data:** Gastroschisis causes significant intestinal dysfunction, with intra-abdominal bowel dilation clinically shown to correlate with worse outcomes. While inflammation and neurovascular impairment have been implicated, genome-wide transcriptional characterization of disease severity remains lacking.

**Methods:** Using a fetal ovine model of complex gastroschisis, in which all gastroschisis animals demonstrated significant intra-abdominal bowel dilation at term, bulk RNA sequencing was performed on proximal small intestinal tissue from mid-gestation and term fetuses across three groups: normal, gastroschisis, and prenatally repaired gastroschisis. Differential gene expression (FDR ≤ .05, |log_2_ fold change| ≥ 1.5) and pathway enrichment analyses were performed, with targeted interrogation of extracellular matrix (ECM), enteric nervous system (ENS), angiogenic, and inflammatory pathways.

**Results:** At mid-gestation, gastroschisis intestine showed minimal transcriptional differences (150 differentially expressed genes [DEGs]) and some bowel dilation. By term, dysregulation was substantial (2,423 DEGs) alongside significant dilation. Normal ontogenetic intestinal maturation patterns were altered, with fewer expected developmental gene changes and discordant pathway regulation. ECM pathway aberrations emerged early and persisted, while ENS, angiogenic, and inflammatory pathways were only dysregulated at term. Fetal repair was associated with normalization of gene expression at term (29 DEGs vs controls).

**Conclusion:** Intestinal transcriptional changes in experimental gastroschisis parallel progressive bowel dilation, consistent with a mechanical stress contribution to intestinal injury. Prenatal repair normalizes both dilation and gene expression, indicating a dynamic and potentially modifiable transcriptional program that supports the rationale for early fetal intervention.

**Mini Abstract:** In a fetal ovine model, progressive bowel dilation in gastroschisis parallels transcriptomic dysregulation of ECM remodeling, neurovascular impairment, and inflammation which is normalized by prenatal repair.

## INTRODUCTION

Gastroschisis is the most common congenital abdominal wall defect, characterized by fetal intestinal herniation into the amniotic cavity. Although postnatal surgical closure is routine, affected infants develop intestinal dysfunction, manifested by delayed enteral feeding and parenteral nutrition dependence. Clinically, gastroschisis is classified as simple or complex. Complex gastroschisis, occurring in 15-20% of cases, is defined by atresia, necrosis, perforation, stenosis, or volvulus and is associated with prolonged intestinal failure, increased healthcare utilization, and higher mortality.^1–3^ Despite these distinctions, intestinal dysfunction remains the central driver of morbidity in both.

Proposed mechanisms of intestinal injury include ischemia from vascular compromise and prolonged exposure of eviscerated bowel to amniotic fluid.^4–6^ These insults promote progressive structural and functional changes including smooth muscle remodeling, fibrosis, and dysmotility.^5^ Intra-abdominal bowel dilation (IABD), particularly prominent in complex disease, correlates with worse clinical outcomes, suggesting that mechanical stress from bowel distension may be associated with additional intestinal injury.^7–11^ However, molecular pathways underlying these changes remain incompletely defined.

The fetal ovine model closely recapitulates human disease following creation of an abdominal wall defect in mid-gestation.^12–14^ We optimized this model to produce a consistent complex gastroschisis phenotype characterized by IABD, impaired motility, and smooth muscle hypertrophy at term.^15, 16^ Prior work demonstrates progressive dilation over gestation in cross-sectional cohorts, consistent with disease evolution, although this remains an indirect inference due to separate animal sampling.^16^ Mechanical stress in other intestinal disease models induces smooth muscle and inflammatory remodeling, supporting a potential role for distension-driven injury.^17–20^ While this model reproducibly recapitulates the bowel dysfunction and structural remodeling seen in human disease, the molecular mechanisms underlying these changes, and whether they are modifiable, remain undefined.

To date, genome-wide transcriptional analyses of gastroschisis, particularly in large animals, are lacking. We therefore performed bulk RNA sequencing of fetal ovine small intestine at mid-gestation and term, with and without fetal repair, to characterize temporal transcriptional programs associated with disease phenotype and to evaluate whether those programs are modifiable with fetal intervention. This hypothesis-generating analysis focused on extracellular matrix (ECM) remodeling, enteric nervous system (ENS), angiogenic, and inflammatory pathways as candidate domains of injury.

## MATERIALS AND METHODS

### Study Design and Experimental Groups

A fetal lamb model of complex gastroschisis was used as previously described.^15, 16^ Gastroschisis was induced at gestational day (GD) 75 (term = 143-145 days). Animals were assigned to: (1) mid-gestation harvest 13-21 days post-induction, (2) term harvest (63-69 days post-induction; GD 138-144), or (3) prenatal repair of the abdominal wall defect at GD 90 (see Supplemental Materials) followed by term harvest (Figure 1A). This yielded five groups: mid-gestation gastroschisis, mid-gestation controls (non-operated littermates), term complex gastroschisis, term controls, and term repaired gastroschisis. Complex gastroschisis was identified by the presence of intestinal stenosis, atresia, volvulus, perforation, or necrosis. Mid-gestation gastroschisis animals demonstrated less IABD at harvest than term gastroschisis animals. All term gastroschisis animals had significant IABD noted prenatally, and prenatally repaired animals demonstrated resolution of IABD at term (Supplemental Figure 1). Procedures were approved by the Ethics Committee on animal experimentation of KU Leuven (057/2021) and conducted in accordance with FELASA (the Federation of European Laboratory Animal Science Associations) guidelines.

**Figure 1.**
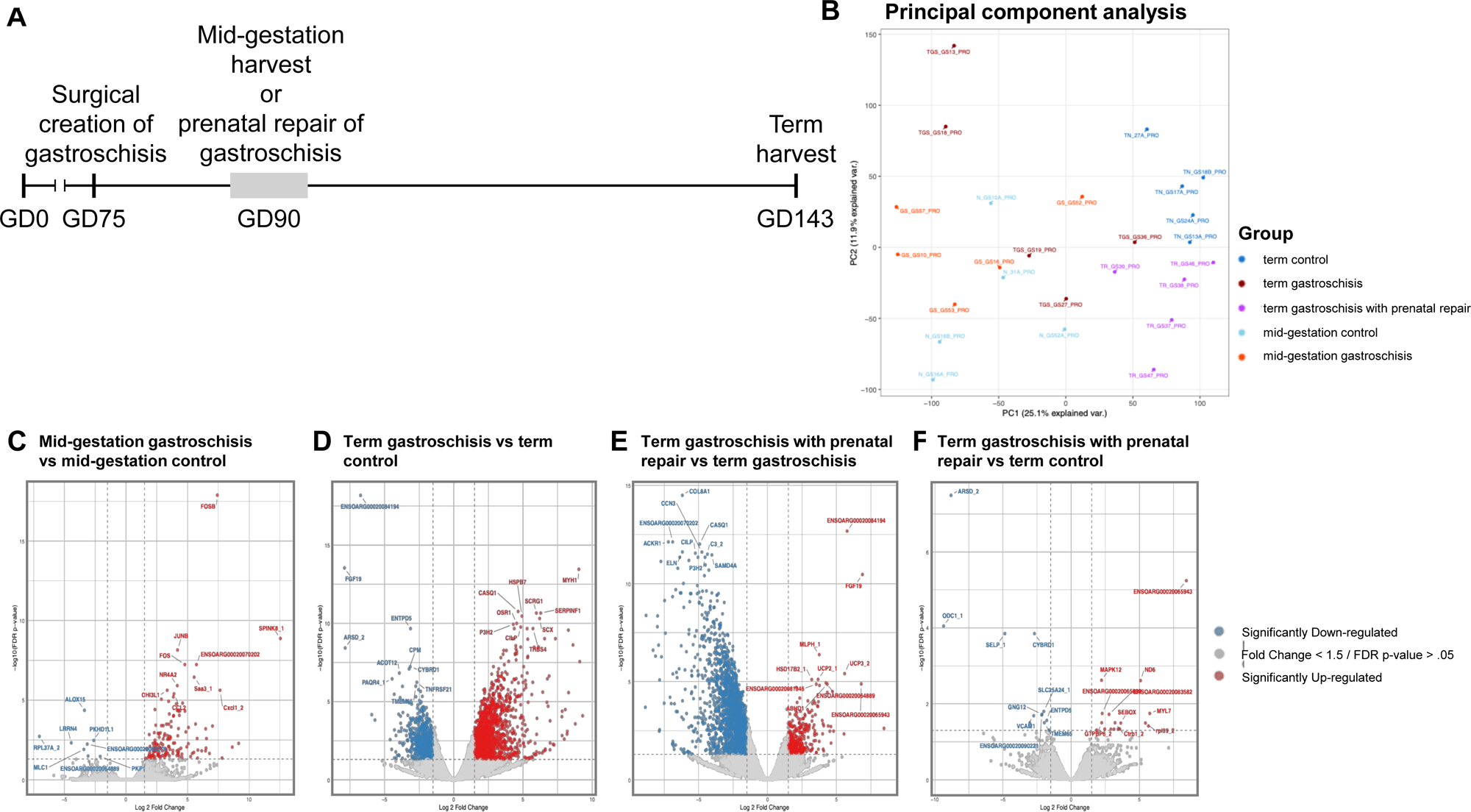
Global transcriptional profiling of the ovine gastroschisis model. (A) Experimental timeline of the ovine gastroschisis model. Gastroschisis was induced at GD75. Mid-gestation fetuses were harvested at GD88–96 (GS n=5; controls n=5). Separate pregnancies progressed to term (GD143) for analysis (term GS n=5; term controls n=5). In a separate cohort, gastroschisis was prenatally surgically repaired via primary closure of the defect at GD90 and harvested at term (repaired n=5). (B) Principal component analysis (PCA) of RNA-seq expression profiles across all groups. (C–F) Volcano plots showing differential gene expression for (C) mid-gestation gastroschisis vs mid-gestation controls, (D) term complex gastroschisis vs term controls, (E) term complex gastroschisis vs term gastroschisis with prenatal repair, and (F) term gastroschisis with prenatal repair vs term control.

### Surgical Model and Tissue Collection

Surgical induction involved creation of a standardized abdominal wall defect with exteriorization of bowel and fixation of a silicone ring for defect stabilization. Fetuses and amniotic fluid were harvested at predefined timepoints. A detailed surgical description and tissue processing methods are provided in the Supplemental Materials, as are histology and immunohistochemistry methods.

### RNA Sequencing and Bioinformatic Analysis

Total RNA was extracted from proximal jejunum. Libraries were prepared using Illumina® Stranded mRNA Library Kit from the University of Houston Seq-N-Edit Core per standard protocols and sequenced using Illumina NovaSeq X+ (∼30 million 2×150 bp reads/sample). Reads were processed and aligned to the ovine genome (ARS-UI_Ramb_v3.0) using CLC Genomics Workbench.^21^

Principal component analysis was performed to assess sample clustering. Differential gene expression was evaluated using a generalized linear model with Benjamini-Hochberg false discovery rate (FDR) correction. Genes with an FDR-*P* value ≤ .05 and |log_2_ fold change| ≥ 1.5 were considered significant. Pathway enrichment analysis was performed using Kyoto Encyclopedia of Genes and Genomes (KEGG) following mapping of ovine genes to human orthologs. Targeted gene set analyses were conducted for ECM, ENS, angiogenic, and inflammatory pathways as follows. A curated ECM gene set was obtained from the MatrisomeDB database (core matrisome and matrisome-associated proteins).^22^ As this type of database was not available for the other pathways, gene sets were derived from relevant KEGG pathways based on biologic plausibility. Specifically, ENS-related genes were obtained from the axon guidance pathway, angiogenesis-related genes from the PI3K-Akt signaling and ECM-receptor interaction pathways, and inflammatory genes from the cytokine-cytokine receptor interaction pathway.

To quantify differences between groups, multivariate analysis of variance (MANOVA) was applied to the first three principal components. Given the small sample size inherent to large animal models, MANOVA results for pathway-specific analyses should be considered exploratory. Models included terms for developmental stage (mid-gestation vs term), disease group (control vs gastroschisis), a separate term indicating repaired status, and an interaction term between the developmental stage and treatment group. Detailed RNA extraction, sequencing, and processing methods are provided in the Supplemental Materials.

### Amniotic Fluid Cytokine Analysis

Inflammatory cytokines were quantified using a multiplex bead-based immunoassay (MILLIPLEX® Ovine Cytokine/Chemokine Panel, SCYT-91K-PX14, Millipore, Burlington, MA) (Supplemental Materials).

### Statistical Analysis

Five animals per group were used for RNA sequencing. This study utilized a subset of animals from a larger, independently generated gastroschisis cohort.^16^ The first five animals from each group meeting inclusion criteria were selected in chronological order of experiment completion. This sample size was optimized for the discovery-based generation of a transcriptomic atlas. Four animals per group (only mid-gestation samples) were used for histologic investigations and at least five samples were used per group (from all five groups) for amniotic fluid cytokine analysis given availability. Statistical analyses were performed using GraphPad Prism v11.0.0 for macOS (GraphPad Software, La Jolla, CA, USA). Data are presented as median with interquartile range and compared using the Mann-Whitney U test for two-group comparisons or the Kruskal-Wallis test with Dunn’s correction for multiple comparisons. *P* < .05 was considered significant.

## RESULTS

### Global Transcriptional Profiling of the Ovine Gastroschisis Model

Principal component analysis (PCA) of proximal jejunum demonstrated clustering by developmental stage and disease state, with mid-gestation samples separating from term along principal component (PC)1 (Figure 1A-B). Gastroschisis samples showed greater variability, while fetal repair samples localized near term controls.

Differential expression was minimal at mid-gestation (150 DEGs) (Figure 1C) but more substantial at term (2,423 DEGs vs control; Figure 1D), paralleling IABD patterns (Supplemental Figure 1). Repaired animals showed minimal differences with controls (29 DEGs; Figure 1F), suggesting attenuation of disease-associated changes at a transcriptional level, while remaining distinct from unrepaired gastroschisis (2,332 DEGs; Figure 1E).

### Gastroschisis Disrupts Gestational Stage-Related Differences of Intestinal Transcriptome

To determine whether gastroschisis alters fetal developmental trajectories, we compared gene expression changes of “normal” control intestinal development between mid-gestation and term with those occurring in gastroschisis animals at those timepoints. Control intestine exhibited 4,370 DEGs, including 1,936 genes upregulated and 2,434 genes downregulated in term compared to mid-gestation controls (Figure 2A).

**Figure 2.**
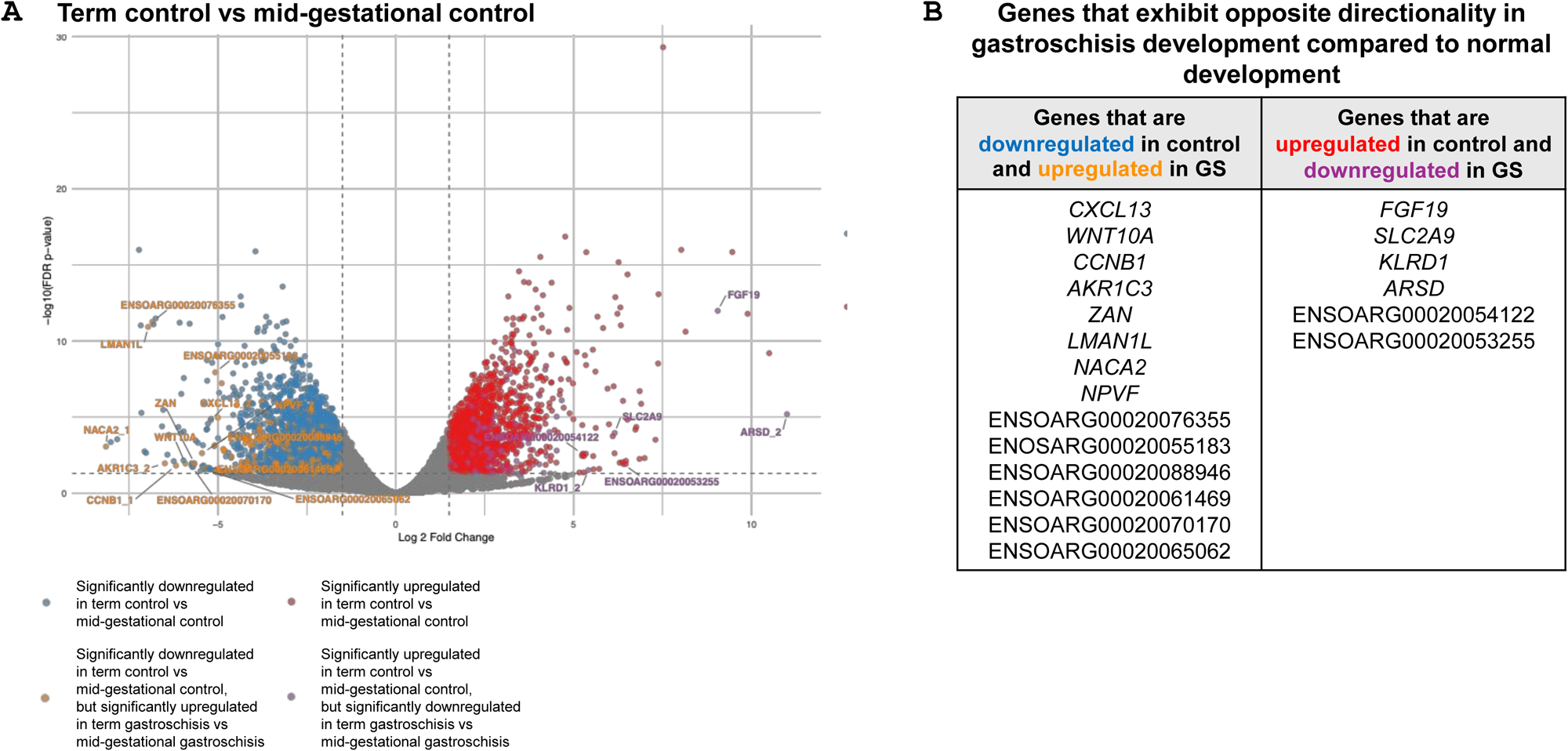
Disruptions in normal intestinal transcriptional maturation in gastroschisis. (A) Volcano plot of differential gene expression between term control and mid-gestation control intestine. Genes significantly upregulated (log₂ fold change ≥ 1.5, FDR ≤ 0.05) are shown in red, and genes significantly downregulated (log₂ fold change ≤ -1.5, FDR ≤ 0.05) are shown in blue. Genes that exhibit opposite directionality in the gastroschisis developmental comparison (term gastroschisis vs. mid-gestation gastroschisis) are highlighted, with normally upregulated genes that become downregulated in gastroschisis shown in purple, and downregulated genes that become upregulated shown in orange. The top 20 genes ranked by absolute log₂ fold change are labeled and (B) organized in a table.

We next examined how these maturation-associated genes behaved in gastroschisis by comparing term to mid-gestation gastroschisis. In contrast to control development, gastroschisis samples exhibited fewer DEGs (539; 304 upregulated, 235 downregulated), suggesting an altered developmental program. While most genes demonstrated consistent directionality between normal development and gastroschisis, a subset demonstrated discordant regulation, with genes that normally increased or decreased during maturation showing the opposite pattern in disease (Figure 2A).

Genes normally downregulated during maturation but remaining elevated in gastroschisis included genes linked to immune chemotaxis (CXCL13), Wnt signaling (WNT10A), cell-cycle progression (CCNB1), and inflammatory steroid metabolism (AKR1C3), suggesting persistent inflammatory, proliferative, and remodeling activity in term gastroschisis intestine. Genes normally upregulated during maturation, including FGF19, SLC2A9, KLRD1, and ARSD that regulate epithelial transport, bile acid signaling, and immune maturation were reduced at term in gastroschisis, suggesting impairments in these functions (Figure 2A-B).

### Pathway-Level Alterations in Gastroschisis

To examine pathways regulated by transcriptional changes in gastroschisis, we performed pathway enrichment analysis followed by focused gene set interrogation in term gastroschisis vs control intestine. KEGG analysis identified enrichment of ECM/mechanotransduction, inflammation, and neurovascular pathways in term gastroschisis (Supplemental Figure 2).

Analysis of selected gene sets revealed patterns distinguishing normal development, gastroschisis, and fetal repair (Figure 3). ECM genes generally demonstrated upregulation in gastroschisis compared to age-matched controls (Figure 3A). This was evident at mid-gestation, more pronounced at term, and normalized following repair, suggesting dynamic profibrotic remodeling that parallels the progression of IABD. To correlate these early transcriptional changes with bowel phenotype, we performed histology on mid-gestation samples, which demonstrated reduced epithelial proliferation (*P* = .03) and non-significant trends toward increased submucosal collagen deposition (*P* = .11) (Supplemental Figure 3).

**Figure 3.**
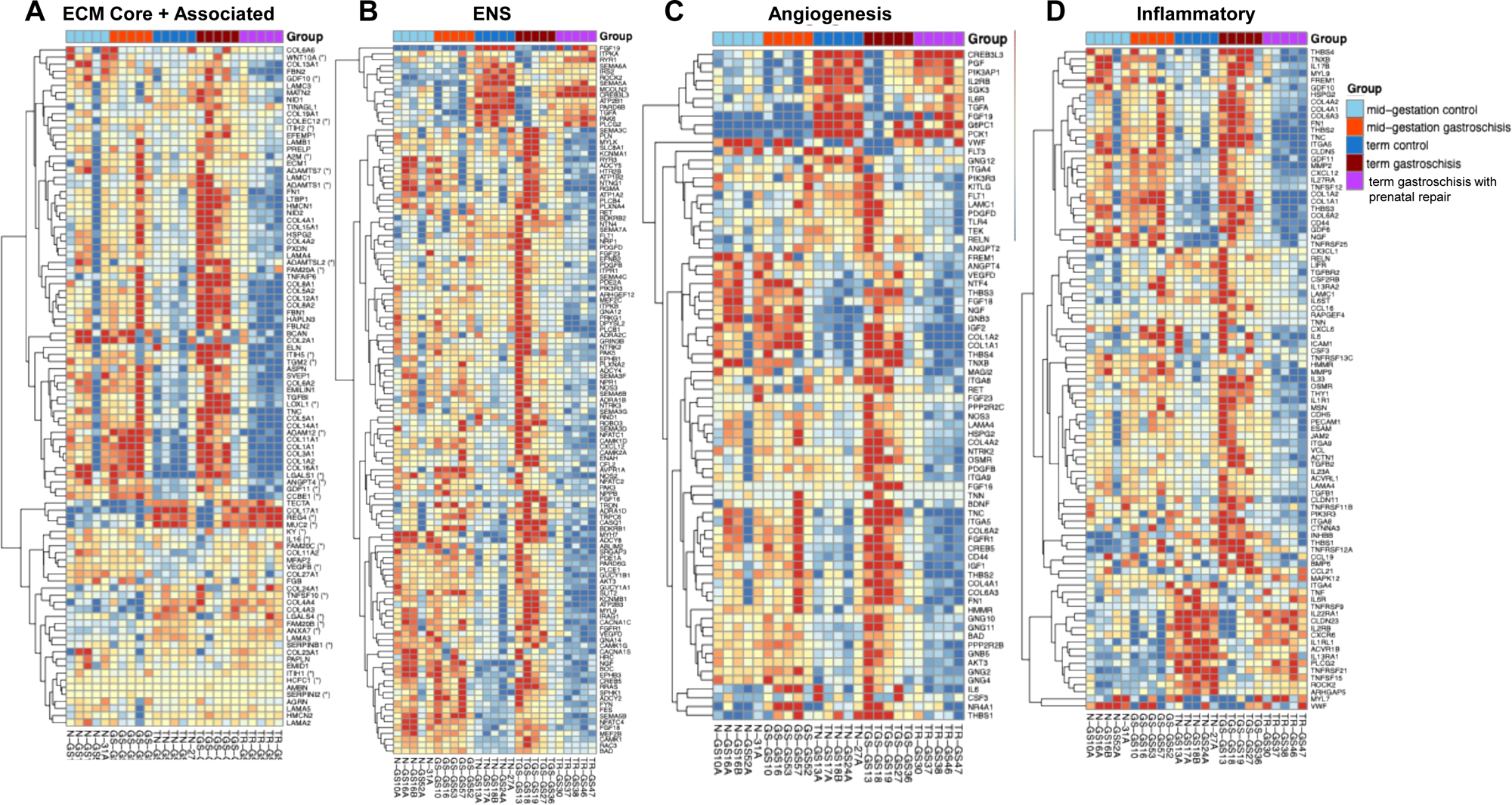
Extracellular matrix, neuro-motility, angiogenesis, and inflammatory pathway remodeling in term gastroschisis small intestine. Heatmaps of differentially expressed (A) ECM genes derived from a curated MatrisomeDB gene set (core matrisome and matrisome-associated proteins), (B) ENS genes contributing to KEGG pathways related to axon guidance, (C) angiogenesis genes related to PI3K-Akt signaling and ECM receptor interaction, and (D) inflammatory genes related to cytokine-cytokine interaction. Rows represent genes and columns represent individual samples, clustered by experimental group. The red-blue color scale on the heatmap indicates the difference x(g, i) - mean(g) between the expression level x(g, i) of gene g in sample i and the mean expression level mean(g) taken over all the samples included in the heatmap. Values above +2 are shown as +2 (indicating 4-fold increased expression over the average for the gene g), while values below -2 are shown as -2 (indicating 4-fold decreased expression below the average for the gene g).

In contrast, ENS, angiogenesis, and inflammatory pathways showed minimal mid-gestation changes but significant dysregulation at term (Figure 3B-D). These patterns were attenuated in the fetal repair group, suggesting modifiability of transcriptional programs. Histology demonstrated early increased ganglionic cellularity (*P* = .03) but no differences in immune cell infiltration (Supplemental Figure 3). These findings support temporal changes leading to intestinal injury in gastroschisis, characterized by ECM remodeling followed by neurovascular and inflammatory dysregulation, with attenuation of these patterns following mid-gestational repair.

### Reversibility of Transcription Changes

Fetal closure of the defect was associated with normalization of gene expression at term; many disease-associated genes shifted toward control levels and reversed directionality (Figure 4A). Notably, the majority of genes differentially expressed in term gastroschisis versus controls demonstrated opposite directionality following repair, with most genes upregulated in gastroschisis becoming downregulated after repair and vice versa.

**Figure 4.**
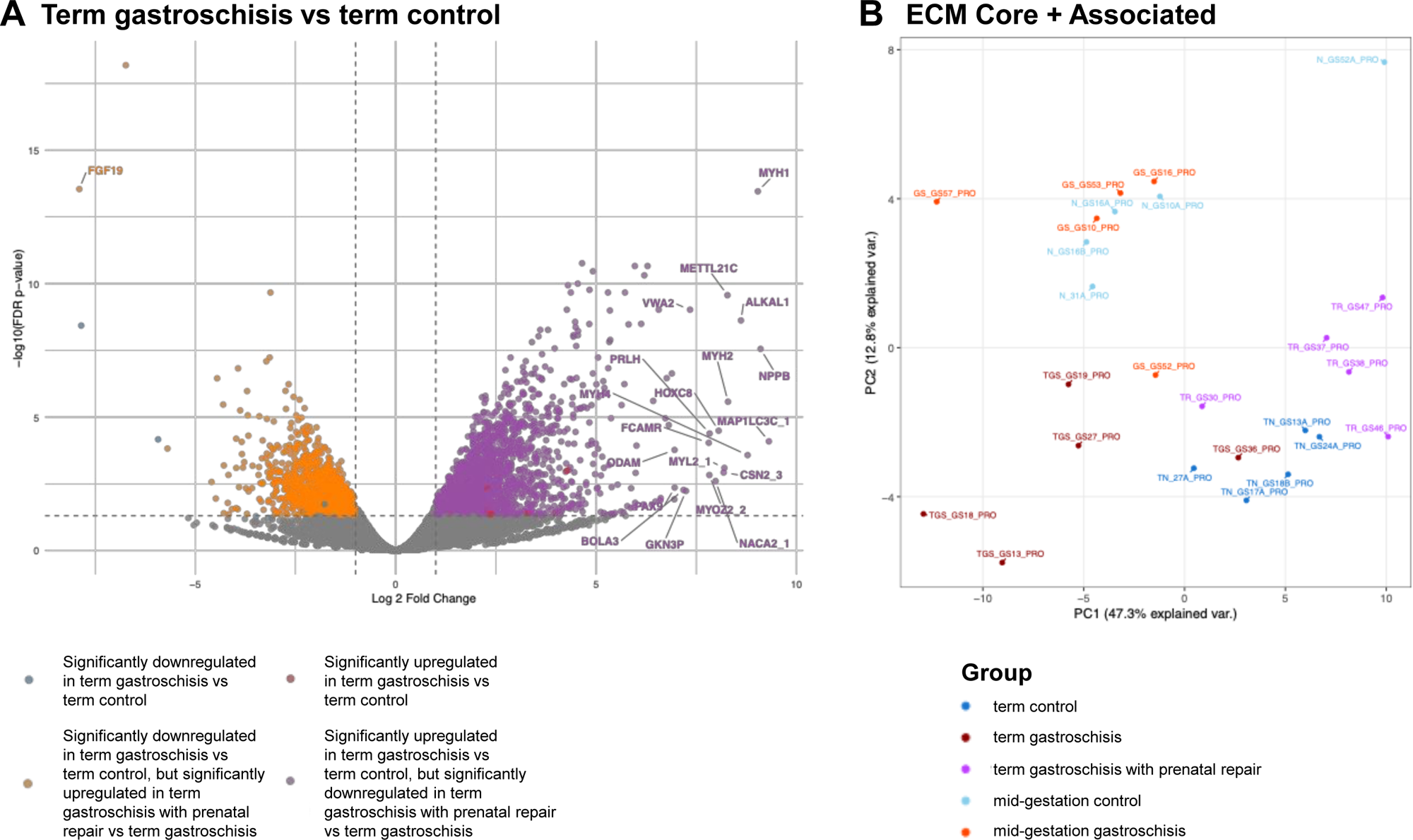
Transcriptional changes with prenatal repair of gastroschisis. (A) Volcano plot of DEGs in term gastroschisis vs term control intestine. Genes significantly upregulated in gastroschisis (log₂ fold change ≥ 1.5, FDR ≤ 0.05) are shown in red, and genes significantly downregulated are shown in blue. Genes demonstrating opposite directionality in the repaired vs unrepaired gastroschisis comparison are highlighted, with upregulated genes that become downregulated after repair shown in purple, and downregulated genes that become upregulated shown in orange. The top 20 genes ranked by absolute log₂ fold change are labeled. (B) PCA of ECM-related genes of interest.

Pathway-specific PCA was performed for ECM, ENS, angiogenic, and inflammatory gene sets (Supplemental Figure 4). ECM genes demonstrated distinct clustering patterns across developmental, disease, and treatment groups (Figure 4B). Term control and repair samples clustered tightly; unrepaired gastroschisis samples were more dispersed. Similar patterns were observed in analyses of inflammatory, angiogenic, and ENS-related gene sets (Supplemental Figure 4). MANOVA showed effects of developmental stage and disease. ECM genes demonstrated significant effects of developmental stage (*P* < .001) and treatment (*P* = .002) without interaction (*P* = .19), consistent with early and persistent dysregulation. In contrast, ENS, angiogenesis, and inflammatory gene sets demonstrated significant stage-dependent interactions (ENS *P* = .02; angiogenesis *P* = .01; inflammatory *P* = .006), indicating transcriptional divergence after the mid-gestation period (Supplemental Figure 4). These findings suggest that mid-gestational repair is associated with attenuation of disease-associated transcriptional programs.

### Amniotic Fluid Cytokine Analysis

Gastroschisis was associated with significantly elevated IL-6, IL-8, IL-10, and VEGF-A at mid-gestation. These levels were reduced following repair and trended toward control values at term, parallelling transcriptional similarities to the control group and suggesting transcriptional programs are reflected in the intrauterine environment (Figure 5).

**Figure 5.**
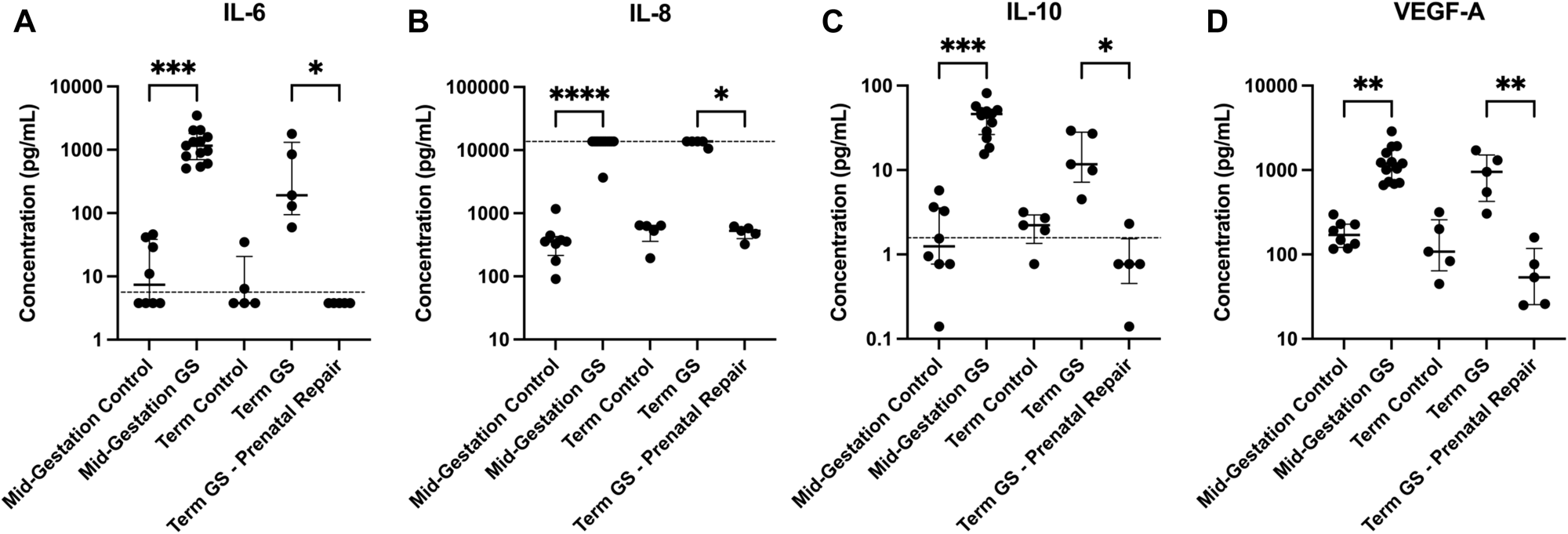
Amniotic fluid cytokine concentrations across gestational and experimental groups. Scatter plots depict concentrations of amniotic fluid (A) IL-6, (B) IL-8, (C) IL-10, and (D) VEGF-A measured by multiplex bead-based immunoassay. (A) IL-6 concentrations were increased in mid-gestation gastroschisis compared with mid-gestation controls (1158 vs 7.412 pg/mL, *P* = .0004) and decreased in repaired gastroschisis compared with unrepaired term gastroschisis (3.795 vs 190.6 pg/mL, *P* = .02). (B) IL-8 concentrations were increased in mid-gestation gastroschisis compared with mid-gestation controls (13,767 vs 355.6 pg/mL, *P* < .0001) and decreased in repaired gastroschisis compared with unrepaired term gastroschisis (527.9 vs 13,767 pg/mL, *P* = .03). (C) IL-10 concentrations were increased in mid-gestation gastroschisis compared with mid-gestation controls (1200 vs 1.245 pg/mL, *P* = .0002) and decreased in repaired gastroschisis compared with unrepaired term gastroschisis (0.768 vs 11.74 pg/mL, *P* = .05). (D) VEGF-A concentrations were increased in mid-gestation gastroschisis compared with mid-gestation controls (1200 vs 170 pg/mL, *P* = .002) and decreased in repaired gastroschisis compared with unrepaired term gastroschisis (53.57 vs 954.8 pg/mL, *P* = .006). Individual points represent biological replicates. Short horizontal bars indicate the median and IQR. Cytokine concentrations are displayed on a log_10_ scale. Dashed horizontal lines denote assay limits of quantification. Statistical comparisons were performed using the Kruskal–Wallis test with Dunn’s multiple comparisons correction. “Term GS – Prenatal Repair” indicates samples from fetuses who underwent repair of the defect at mid-gestation and were harvested at term.

## DISCUSSION

Gastroschisis is associated with persistent intestinal dysfunction, yet molecular programs underlying this phenotype remain incompletely defined. This study characterizes the transcriptomic landscape of the developing intestine in a fetal ovine model and demonstrates that these changes are partially modifiable following fetal repair. We identify early ECM-related changes followed by later alterations in selected pathways related to neuro-motility, vascular, and inflammatory signaling. The number of DEGs paralleled the extent of IABD across groups: more substantial at term than mid-gestation, and normalized following repair. While consistent with a model in which dilation-associated mechanical stress contributes to intestinal injury, this association is correlative as the design cannot exclude alternate upstream contributors. KEGG analysis identified candidate pathways that may contribute to intestinal dysfunction, and the transcriptional response appears dynamic and potentially modifiable.

Gastroschisis was associated with activation of pathologic remodeling pathways and disruption of changes seen between the normal intestinal tissue at mid-gestation and term timepoints. Genes that normally declined at term were paradoxically sustained, including inflammatory mediators (e.g. CXCL13) and regulators of proliferation and signaling (e.g. WNT10A, CCNB1), while key epithelial maturation genes (e.g. FGF19) did not increase as expected at term. These trajectory-switching patterns suggest that exposed bowel sustains an immature, pro-inflammatory transcriptional state, consistent with prior reports of delayed epithelial maturation and impaired absorptive function in gastroschisis.^23, 24^

A central finding of this study is the early and progressive association of ECM pathway dysregulation. Transcriptomic pathway analyses demonstrated coordinated upregulation of genes involved in matrix organization, focal adhesion, integrin signaling, and TGF-β–associated pathways, consistent with a profibrotic remodeling program. These findings align with prior descriptions of smooth muscle hypertrophy in ovine and human gastroschisis.^25, 26^ Similar responses are observed in models of intestinal mechanical stress, including distraction enterogenesis and bowel obstruction, where mechanical stretch induces ECM deposition and smooth muscle proliferation in human and animal models.^19, 27, 28^ IABD is a consistent feature of both the ovine model and human gastroschisis, and could represent an upstream driver of injury, though this model cannot separate the effects of mechanical stress from other injury mechanisms, such as amniotic fluid exposure or vascular compromise.^16^ Nonetheless, the data support a plausible explanation for complex gastroschisis pathophysiology, in which progressive fetal bowel dilation generates mechanical stress that triggers ECM remodeling and smooth muscle hypertrophy, establishing structural alterations leading to intestinal dysfunction. This process may, at least in part, represent the second “hit” in the pathogenesis of gastroschisis, hypothesized to result from an initial defect due to rupture of the physiological umbilical hernia followed by progressive injury to the eviscerated bowel.^4^

In contrast to early and persistent ECM changes, disruption of selected neuro-motility, vascular, and inflammatory signaling pathways emerged later in the cross-sectional comparison, suggesting these alterations may represent secondary responses to ongoing remodeling. ENS-related genes in the axon-guidance KEGG pathway demonstrated dysregulated expression at term, consistent with impaired neuromuscular development and aligning with known abnormalities in the ENS in gastroschisis.^29, 30^ Angiogenic pathways (PI3K-Akt signaling and ECM receptor interaction) also exhibited heterogeneous dysregulation, potentially related to mesenteric constriction or altered intrauterine perfusion.^31^ Many inflammatory genes (cytokine-cytokine interaction KEGG pathway) were upregulated in gastroschisis, while a subset induced during normal intestinal maturation—including a marker of immune surveillance, KLRD1—failed to increase as expected. Gastroschisis may involve both pathologic inflammatory activation and failure to engage normal developmental immune programs, consistent with reports of premature T cell activation and aberrant ILC2 expansion in exteriorized gastroschisis bowel.^23^ The presence of persistent chemokine signaling (CXCL13) and altered metabolic regulators (AKR1C3) further supports a model of dysregulated immunity. These findings were mirrored by alterations in the intrauterine environment. Amniotic fluid contained elevated inflammatory and angiogenic mediators at mid-gestation. Levels were significantly reduced in the repair cohort, supporting the dynamic nature of the disease. Notably, elevated IL-6 and IL-8 levels in gastroschisis amniotic fluid are consistent with prior reports.^26^

A novel insight is that transcriptional changes appear to be modifiable in lambs undergoing mid-gestational primary closure. Most genes dysregulated in gastroschisis demonstrated opposite directionality in the repair cohort, suggesting restoration toward a normal trajectory. This was observed across ECM, inflammatory, and neurovascular pathways, suggesting intestinal dysfunction represents a modifiable state rather than irreversible injury. However, whether it reflects reversal of established injury or prevention of further progression cannot be determined from this study design, as repair occurred before the period of maximal transcriptional differences.

These findings are consistent with experimental models demonstrating that removal of mechanical stress can reverse structural and functional changes in the intestine, including smooth contractility.^28^ From a developmental perspective, this suggests the existence of a window during which interruption of pathologic stimuli permits re-engagement of normal maturation programs. The observation that ECM remodeling occurs early further suggests that intervention timing may be critical to prevent disease progression.

This study has several limitations. First, all comparisons are cross-sectional across independent animal cohorts, preventing inference about temporal progression within individuals. Thus, gene expression changes and amniotic fluid cytokine levels represent between-group differences but do not directly demonstrate progressive disease within individuals. As a hypothesis-generating analysis, the transcriptomic findings identify associations rather than establish causality, and additional studies are needed to link changes to downstream protein-level or functional effects. Importantly, because all surviving term animals in this model exhibited complex gastroschisis, we were unable to compare simple and complex phenotypes directly; while this reflects the robustness and consistency of the model, it limits our ability to identify phenotype-specific transcriptional mechanisms.

The small number of biological replicates, inherent to large animal models, limits statistical power, increases the contribution of between-animal variability, and may affect the reliability of multivariate analyses applied to pathway-specific gene sets. In our analyses, DEGs were defined using FDR < .05. Because of the limited sample size, we also applied a stringent fold-change threshold (|log₂FC| ≥ 1.5) to prioritize genes with larger magnitude changes, recognizing that this approach may exclude biologically relevant genes with smaller effect sizes.

Our analysis focused on predefined gene sets, including ECM genes from the Matrisome database, axon guidance genes (KEGG), angiogenic pathways (KEGG PI3K-Akt signaling and ECM-receptor interaction), and inflammatory pathways (KEGG cytokine-cytokine interaction), guided by prior literature implicating these domains in gastroschisis-associated intestinal dysfunction and by their relevance to mechanotransduction and bowel wall remodeling.^26, 29, 32–34^ However, this targeted approach introduces selection bias, as transcriptomic analyses may overlook other relevant pathways. Statements regarding the relative timing of pathway activation may reflect the composition of the selected gene sets rather than true biological sequencing. Accordingly, our findings should be interpreted as prioritizing plausible candidates rather than providing comprehensive unbiased discovery.

While the ovine model closely recapitulates human gastroschisis, species-specific differences in gene annotation and developmental timing may limit direct translation of findings. Cross-species mapping of ovine genes to human orthologs captures a large majority of ovine protein-coding genes, but genes lacking high-confidence orthologs or clear one-to-one relationships were excluded, which can bias enrichment analyses toward evolutionarily conserved pathways and core mammalian functions.^35^ Finally, although switched transcriptional directionality following prenatal repair of gastroschisis suggests potential modifiability of some molecular features of injury, the extent to which this reflects reversal versus interruption of ongoing disease progression remains uncertain. Because this was a surgically induced model rather than an embryonic model of disease initiation, the developmental timing of intervention cannot be directly mapped to human gestation, and it is not possible to define what would constitute “early” intervention in a clinical setting. Moreover, whether fetal repair can prevent progression to more complex intestinal pathology remains unknown, as disease evolution may still occur despite prenatal correction. Nonetheless, these findings support the broader concept that the intrauterine environment contributes to the intestinal phenotype and may represent a potential window for therapeutic modulation.

In summary, gastroschisis in the fetal lamb model induces coordinated, temporal, and modifiable transcriptomic changes, characterized by early ECM remodeling followed by neurovascular and inflammatory dysregulation. The changes in DEGs associated with the transcriptional changes parallel the changes in IABD. These findings support the development of future studies to test the hypothesis that dilation-associated mechanical stress contributes to intestinal injury which is dynamic and modifiable by early intervention. By establishing the transcriptional changes related to gastroschisis-associated intestinal dysfunction, this study provides a framework for future mechanistic studies and informs the rationale for prenatal therapeutic strategies.

## Abbreviations

CV: coefficient of variation
DEG: differentially expressed genes
ECM: extracellular matrix
ENS: enteric nervous system
FDR: false discovery rate
FELASA: Federation of European Laboratory Animal Science Associations
GD: gestational day
GO: gene ontology
IABD: intra-abdominal bowel dilation
IFN: interferon
IHC: immunohistochemistry
IL: interleukin
IP: interferon gamma-induced protein
KEGG: Kyoto Encyclopedia of Genes and Genomes
LLOQ: lower limit of quantification
MIP: macrophage inflammatory protein
PBS: phosphate-buffered saline
PC: principal component
PCA: principal component analysis
PFA: paraformaldehyde
ROI: region of interest
RPKM: reads per kilobase of transcript per million mapped reads
SYP: synaptophysin
TMM: trimmed mean of M-values
TNF: tumor necrosis factor
TOM: topological overlap matrices
ULOQ: upper limit of quantification
VEGF: vascular endothelial growth factor

## Acknowledgments

The authors would like to thank Cecilia Ljungberg, PhD and the RNA In Situ Hybridization Core facility at Baylor College of Medicine and Texas Children’s Hospital for assistance with slide imaging using the Zeiss Axioscan.Z1. We also thank Shixia Huang, PhD and the Proteomics Core at Dan L. Duncan Cancer Center at Baylor College of Medicine for assistance with amniotic fluid multiplex cytokine analysis. Dennis Wylie, PhD and Dhivya Arasappan, MS of the Bioinformatics Consulting Group at the University of Texas at Austin were instrumental in providing technical assistance with bioinformatics analysis. We are also grateful to Keely Wolf, MS in the Department of Surgery at Baylor College of Medicine for biostatistics consulting.

## FIGURE LEGENDS

**Supplemental Figure 1**. MRI-derived intra-abdominal bowel dilation. (A) Differences in mid-gestation IABD between control (n = 1) and gastroschisis (n = 3). (B) At term, MRI-IABD was significantly increased in gastroschisis compared with controls (22.3 mm vs 5.3 mm, *P* = .03). Prenatal repair normalized bowel dilation, with no difference between repaired gastroschisis and controls (5.7 mm vs 5.3 mm, *P* > .99), and significantly reduced dilation compared with unrepaired gastroschisis (5.7 mm vs 22.3 mm, *P* = .02)(n = 5 per group). Data are presented as median with interquartile range. Statistical comparisons for (B) were performed using Dunn’s multiple comparisons test. “Term GS – Prenatal Repair” indicates samples from fetuses who underwent repair of the defect at mid-gestation and were harvested at term.

**Supplemental Figure 2:** KEGG pathway enrichment analysis of differentially expressed genes (DEGs) in term gastroschisis versus term control small intestine demonstrating pathways clustering into ECM and mechanotransduction (e.g., ECM-receptor interaction, focal adhesion, integrin signaling), inflammatory signaling (e.g., cytokine-cytokine receptor interaction, TGF-β signaling), and neurovascular signaling (e.g., axon guidance, calcium signaling, cGMP-PKG signaling). Dot plot displays the top enriched pathways ranked by adjusted *P* value (FDR < 0.05). Dot size represents gene count and color intensity corresponds to −log10(adjusted *P* value). Arrows denote pathways selected for focused gene set interrogation.

**Supplemental Figure 3.** Histologic and immunohistochemical characterization of proximal jejunum in mid-gestation control and gastroschisis lambs. (A-B) Representative images of proximal jejunal crypt-villus units stained for CD45 (magenta) in control (A) and gastroschisis (B) lambs. (C) Quantification of CD45+ cells in crypt-villus units and submucosa demonstrated no significant differences between groups (crypt-villus units: 15.47% vs 18.63%, *P* = .89; submucosa: 5.51% vs 6.52%, *P* = .69). (D-E) Representative images of intestinal crypts stained for Ki67 (magenta) in control (D) and gastroschisis (E) lambs. (F) Gastroschisis animals demonstrate reduced crypt proliferation, as evidenced by decreased Ki67+ cells (41.91% vs 31.62%, *P* = .03). (G-H) Representative images of enteric ganglia stained for synaptophysin (SYP; magenta) in control (G) and gastroschisis (H) lambs. (I) Gastroschisis lambs demonstrated increased cells per ganglion (12.85 vs 18.65, *P =* .03). (J-K) Representative trichrome-stained sections in control (J) and gastroschisis (K) animals, with collagen highlighted in blue. (L) Slides appeared to have increased submucosal collagen deposition in gastroschisis, but quantification of mean collagen intensity did not reach statistical significance (0.2075 vs 0.4156, *P* = .11). Data are presented as median with interquartile range; n = 4 per group. Statistical comparisons were performed using the Mann-Whitney U test. Scale bars: 100 um (A-B), 20 um (D-E, G-H), and 50 um (J-K).

**Supplemental Figure 4.** Pathway-specific PCA for (A) ECM, (B) inflammatory, (C) angiogenic, and (D) ENS gene sets. Arrows indicate the direction and magnitude of coefficient vectors fit by a one-way MANOVA model based on PCs 1-2 in which the “term control” group is taken as the reference group, thus fitting a 2-dimensional (PCs 1-2) coefficient vector representing the expected difference in PC position of samples from each of the other 4 groups (4 vectors shown in total). (E) MANOVA showing effects of gestational timepoint and disease. Models included terms for gestational timepoint, disease group (control vs. gastroschisis), a separate term indicating repaired status, and an interaction term between developmental stage and disease group.

