## Supplemental Figures 1-4 for "Intra-Abdominal Bowel Dilation in Experimental Gastroschisis is Associated with a Modifiable Transcriptomic Program of Intestinal Dysfunction"

Supplemental Figure 1

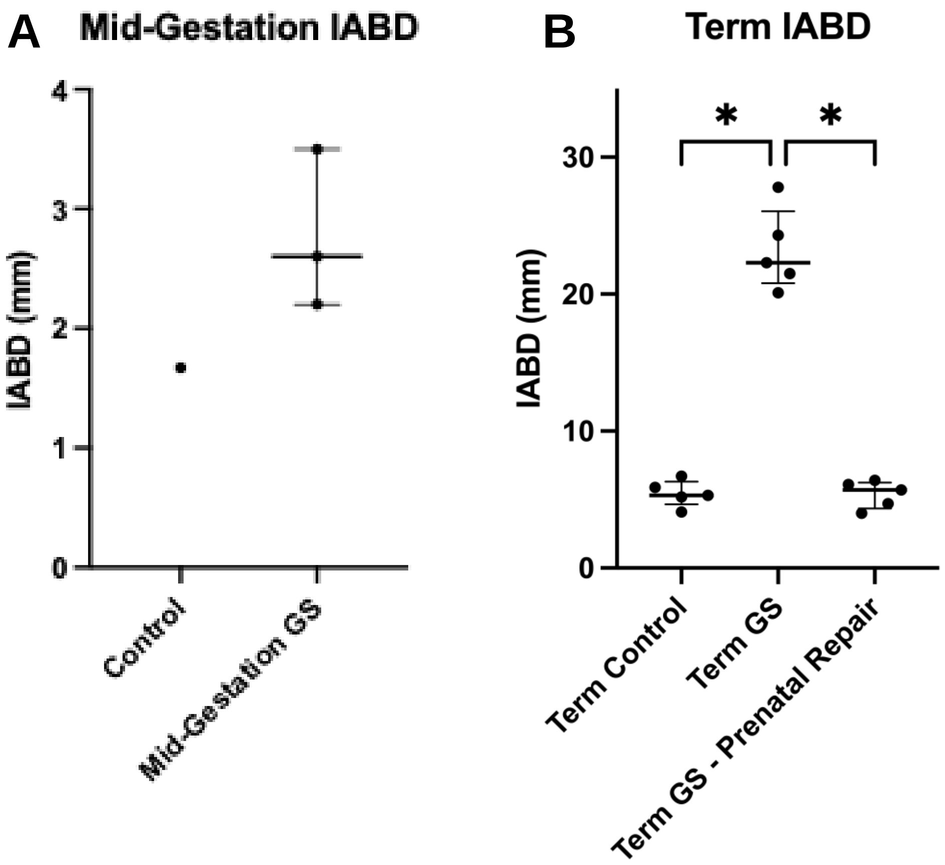

Supplemental Figure 2

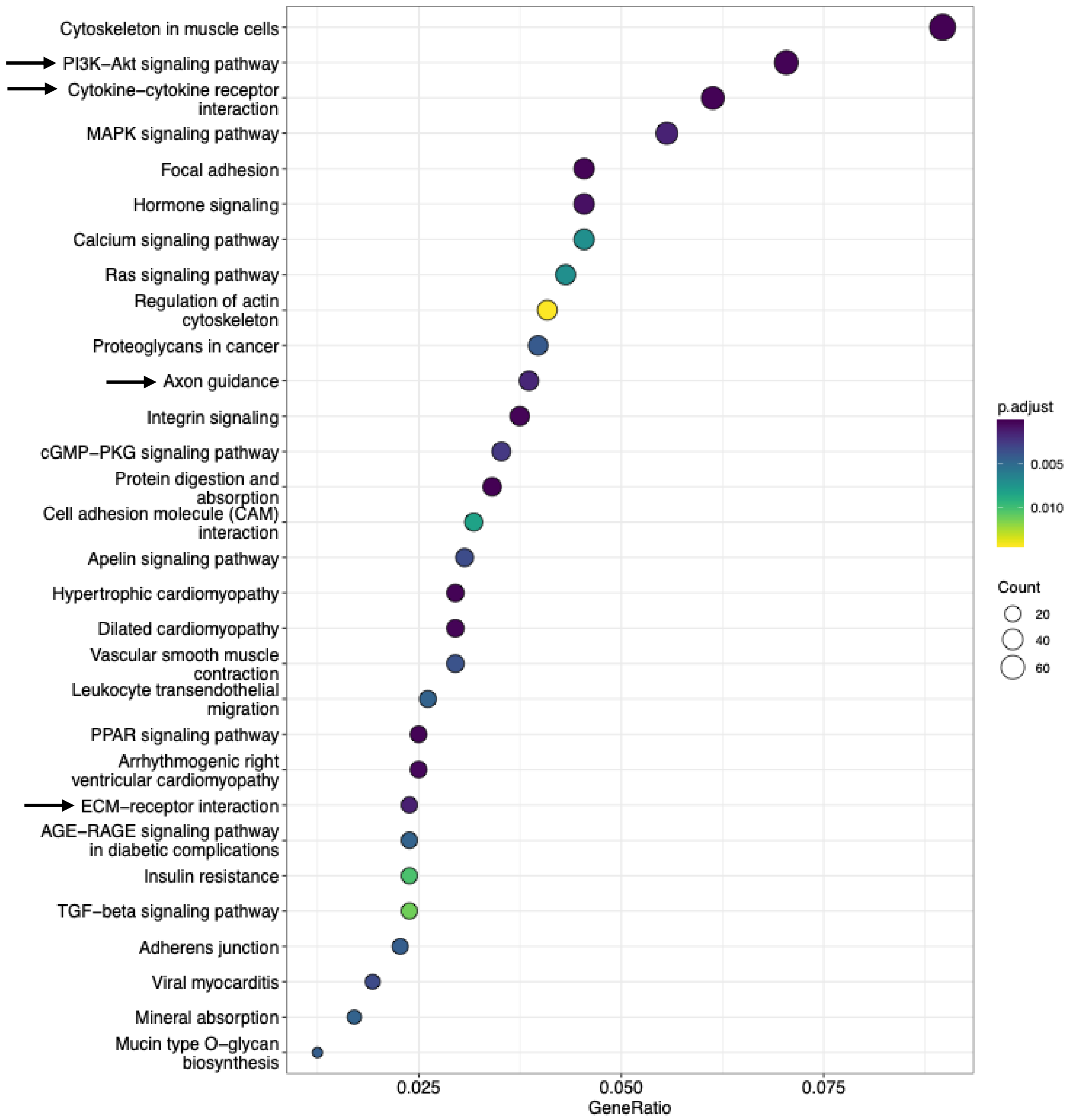

Supplemental Figure 3

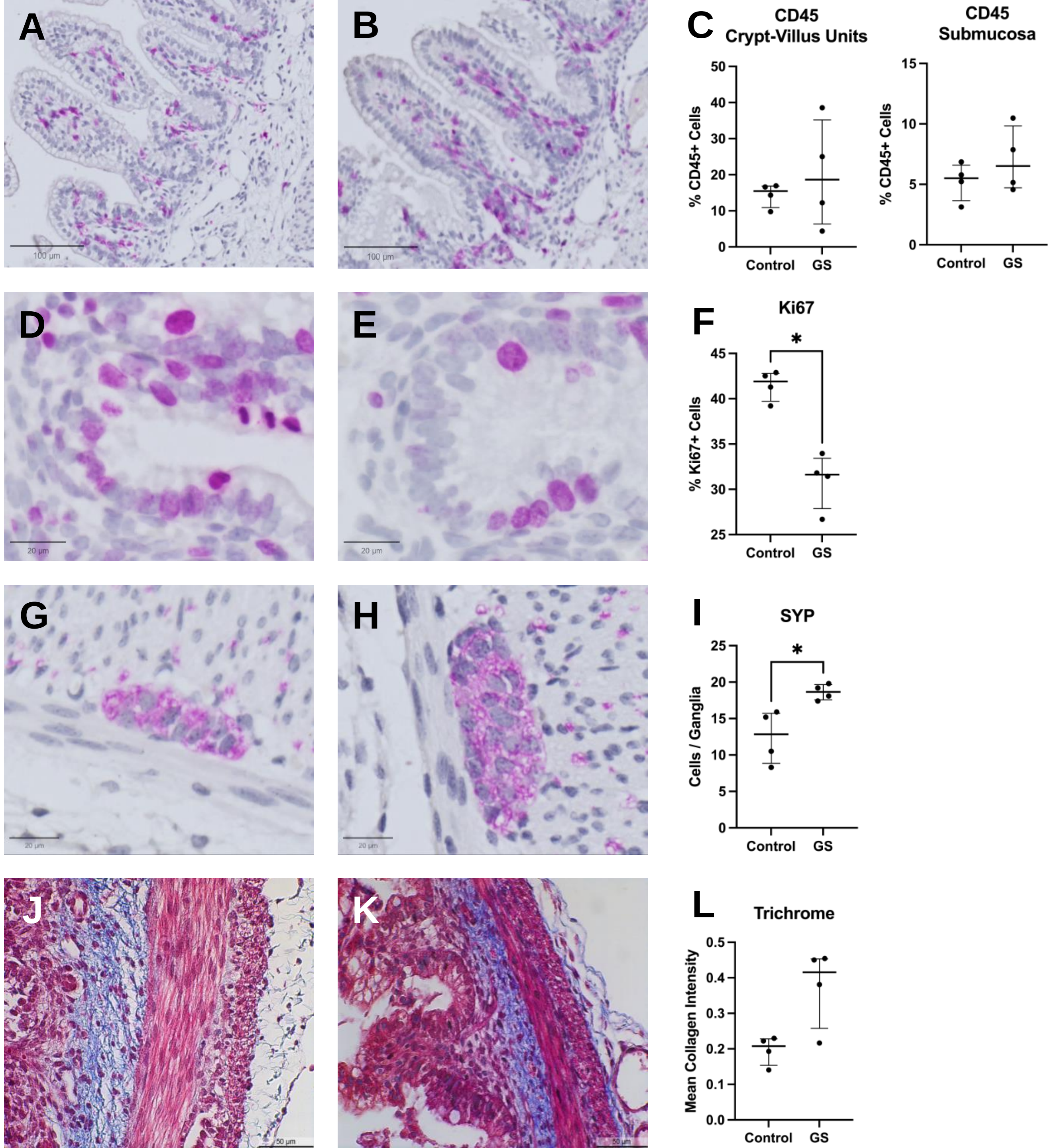

Supplemental Figure 4

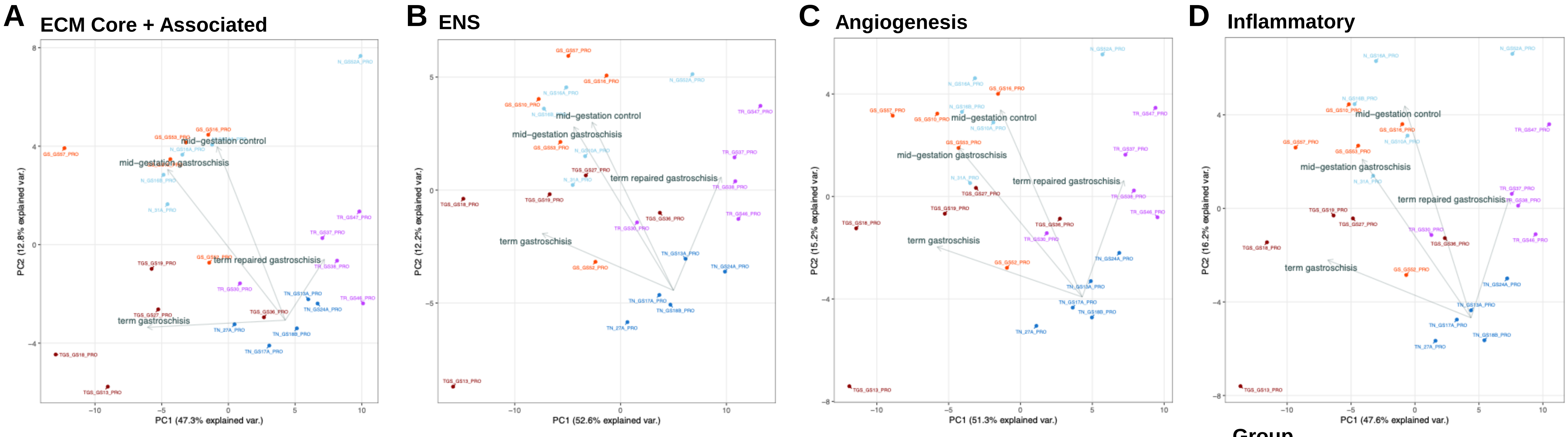


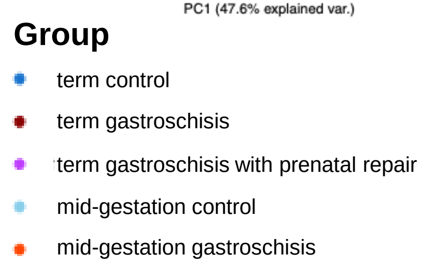


**E**

| Gene Set | Term *P* | Disease *P* | Interaction *P* |
|---|---|---|---|
| ECM (combined) | < .00001 | .002 | .19 |
| ENS | .00002 | .001 | .02 |
| Angiogenesis | < .00001 | .0006 | .01 |
| Inflammatory | < .00001 | .0006 | .006 |
