## Supplemental Methods for "Intra-Abdominal Bowel Dilation in Experimental Gastroschisis is Associated with a Modifiable Transcriptomic Program of Intestinal Dysfunction"

#### Surgical Model and Tissue Collection

Under maternal general anesthesia, the fetal abdomen was exposed via maternal midline laparotomy and hysterotomy. A standardized full-thickness abdominal wall defect was created in the fetus, a 1 cm diameter silicone ring was secured to the wall for stabilization of the defect, and the bowel from proximal jejunum to distal spiral colon was exteriorized. The fetus was returned to the uterus and the uterus and maternal abdomen were closed. Fetuses were harvested and their tissue collected (bowel and amniotic fluid) at predetermined gestational time points. Detailed characterization of the model, including the development of complex gastroschisis and associated intestinal injury, has been previously reported.[1]

#### Fetal Repair of Surgically Induced Gastroschisis

Fetal repair was performed at mid-gestation (GD 85–90) under maternal general anesthesia via laparotomy and hysterotomy. The fetal abdomen and hindlimbs were exteriorized, and the silicone ring was removed. The mesenteric stalk was dissected from the abdominal wall defect, which was then enlarged to approximately 4 cm to facilitate reduction of the exteriorized bowel into the abdominal cavity. The defect was subsequently closed primarily. The fetus was returned to the uterus, and the pregnancy was continued until term.

#### Tissue Collection

At the designated gestational time point, fetuses were delivered by cesarean section under maternal anesthesia. Amniotic fluid samples were taken at the time of harvest via hysterotomy, centrifuged at 3000 rpm for 10 minutes immediately after collection, and the supernatant was stored at -80 °C. Following euthanasia of the ewes and fetuses, intestinal tissue was harvested for molecular and histologic analyses. Segments of proximal jejunum, identified based on the presence of arterial supply from the first two branches of the cranial mesenteric artery, were rapidly dissected and rinsed in sterile phosphate-buffered saline (PBS). Intestinal tissue to be used for RNA sequencing was frozen and stored at -80 °C. Bowel specimens to be used for immunohistochemical analysis were immersed in 4% paraformaldehyde (PFA) for 24 hours, washed with PBS for one hour, and transferred to 70% ethanol for dehydration prior to being embedded in paraffin longitudinally and transversely.

**RNA Extraction, Library Preparation, and Sequencing**

Frozen bowel samples were lysed and homogenized, and RNA was extracted using a PureLink™ RNA Mini Kit (Invitrogen, Carlsbad, CA, USA) following the manufacturer’s protocol. Extracted RNA underwent quantification using Qubit Fluorometer High Sensitivity RNA Assay (Thermo Fisher Scientific, Waltham, MA, USA). Five hundred ng of the extracted RNA was used for the library preparations and sequenced at the University of Houston Seq-N-Edit Core per standard protocols. RNA was enriched and mRNA libraries were prepared with Illumina® Stranded mRNA Library Kit (LC Sciences, Houston, TX, USA). The size selection for libraries was performed using SPRIselect beads (Beckman Coulter, Brea, CA, USA) and purity of the libraries was analyzed using the High Sensitivity d1000 ScreenTape using Agilent Tapestation 4200. The prepared libraries were pooled and sequenced using Illumina NovaSeq X+, generating ~30 million 2x150 bp paired end reads per sample.

**RNA Data Processing and Analysis**

Raw sequencing data (FASTQ files) were processed using CLC Genomics Workbench (Version 25.0.3, Qiagen, Germantown, MD, USA). Adapter sequences were trimmed and reads were mapped to the sheep reference genome (Ovis aries, ARS-UI_Ramb_v3.0; Ensembl release 115; assembly accession GCA_016772045.2). Alignment parameters included mismatch cost 2, insertion cost 3, deletion cost 3, length fraction 0.8, similarity fraction 0.8, and a maximum of 10 mapping locations per read. Integer read counts were normalized using the trimmed mean of M-values (TMM) algorithm. Principal component analysis was performed to assess sample variability and clustering. To quantify differences between groups, multivariate analysis of variance (MANOVA) was applied to the first three principal components. Models included terms for developmental stage (mid-gestation vs term), disease group (control vs gastroschisis), a separate term indicating repaired status, and an interaction term between the developmental stage and treatment group.

Differential gene expression was evaluated using a generalized linear model (GLM) framework. P-values were adjusted for multiple comparisons using the Benjamini-Hochberg false discovery rate (FDR). Genes with an absolute $\log_2$ fold change (LFC) greater than 1.5 and an FDR-adjusted $P$ value < .05 were considered significantly differentially expressed. Kyoto Encyclopedia of Genes and Genomes (KEGG) pathway enrichment analysis was performed using clusterProfiler package after mapping ovine genes to human orthologs through Ensembl BioMart, retaining only high-confidence one-to-one orthologs. KEGG pathway enrichment was conducted against the human KEGG database. A curated ECM gene set was obtained from the

MatrisomeDB database (core matrisome and matrisome-associated proteins) and used for focused ECM pathway analysis.[2] As this type of database was not available for the other pathways of interest, gene sets were derived from relevant KEGG pathways based on biologic plausibility. Specifically, ENS-related genes were obtained from the axon guidance pathway, angiogenesis-related genes from the PI3K-Akt signaling and ECM-receptor interaction pathways, and inflammatory genes from the cytokine-cytokine receptor interaction pathway.

**Histology and Immunohistochemistry**

Paraffin embedded sections were cut at 5 µm using a Leica RM2155 automatic microtome (Leica Biosystems, Wetzlar, Germany) and were mounted on slides. Slides were deparaffinized and rehydrated to PBS following standard protocol. A standard immuno-histochemistry (IHC) protocol was followed for all stains. Primary antibodies included mouse anti-CD45 for inflammatory cells (Clone 1.11.32, BioRad, 1:200), mouse anti-synaptophysin (SYP) for enteroendocrine cells (Clone DAK-SYNAP, Dako, 1:1000), and mouse anti-Ki67 (Dako, Clone MIB-1, 1:200). Antibodies were detected by EnVision FLEX HRP Magenta Substrate Chromogen System Kits (Dako North America, Carpinteria, CA) and counterstained with hematoxylin. Immunohistochemistry slides were digitized using Zeiss Axioscan.Z1 with 20X/0.8 NA lens and Zeiss Zen 3.9 software.

Trichrome staining was performed manually using a Trichrome, Masson, Aniline Blue Stain Kit (Newcomer Supply, Waunakee, WI, USA) following the manufacturer’s protocol. Trichrome slides were imaged with Leica DM 2000® using Leica Application Suite X® version 3.0.4.16529.

### Histologic Quantification

Whole-slide images were analyzed using QuPath (version 0.6.0). Regions of interest (ROI) were manually annotated for each target protein. ROIs for the CD45-stained tissue sections were chosen from longitudinal sections and defined as well-defined crypt-villus units and regions of the submucosa underlying the crypt-villus units.[3] ROIs for the Ki67-stained tissue sections were limited to well-defined crypts of longitudinal sections.[3] Twenty ROIs were chosen for each sample. Automated cell detection was performed using the built-in nuclear detection algorithm to identify individual cells after color deconvolution and manual adjustment of stain vectors for clear separation between the magenta stain and hematoxylin. The percentage of positively stained cells was calculated for each region and exported for statistical analysis.

Enteroendocrine cell ganglia were identified in longitudinal tissue sections based on SYP staining bundles. Ten random well-defined ganglia were chosen for each sample. The nuclei in each ganglion were manually counted, and the cell counts were exported for statistical analysis.

Collagen content was quantified using trichrome-stained transverse tissue sections. Color deconvolution was performed with manual adjustment of the stain vectors to ensure adequate separation. The mean pixel intensity of the blue channel in the submucosal layer was measured and exported for statistical analysis.

### Amniotic Fluid Cytokine Analysis

Amniotic fluid samples were thawed on ice, centrifuged at 14,000 × g at 4 °C for 5 minutes, and the supernatant was analyzed in duplicate (25 µl per well). Analytes measured included interferon (IFN) γ, interleukin (IL)-1α, IL-1β, IL-4, IL-6, IL-8 (CXCL8), IL-10, IL-17A, IL-36RA, interferon gamma-induced protein 10 (IP 10) (CXCL10), macrophage inflammatory protein (MIP)-1α (CCL3), MIP-1β (CCL4), tumor necrosis factor (TNF)-α, and vascular endothelial growth factor (VEGF)-A. Samples were incubated with antibody-coupled beads overnight (16-18 hours, at 4 °C) followed by biotinylated detection antibodies and streptavidin-phycoerythrin. Data were acquired on a Bio-Plex 200 system (Bio-Rad, Hercules, CA) with ≥ 50 beads collected per analyte and concentrations calculated using a five-parameter logistic standard curve in Bio-Plex manager software (v7.2). Replicate values were averaged for analysis. Measurements were reviewed for assay quality based on bead counts and replicate coefficients of variation (CV). Values below the lower limit of quantification (LLOQ) were assigned a value of LLOQ/2 for statistical analysis. Values exceeding the upper limit of quantification (ULOQ) were assigned the ULOQ value.[4] In instances where concentrations fell below the lower standard curve range but remained within the measurable dynamic range of the assay, concentrations were extrapolated using the fitted regression model generated by the five-parameter logistic curve.
